# Cell signaling pathways in human mutant PAX6 corneal cells: an in vitro model for aniridia-related keratopathy

**DOI:** 10.1101/2021.03.19.436143

**Authors:** Marta Słoniecka, André Vicente, Berit Byström, Fátima Pedrosa Domellöf

## Abstract

**PURPOSE:** To establish an *in vitro* model of aniridia-related keratopathy (ARK) using CRISPR/Cas9 engineered human keratocytes with mutations in the PAX6 gene, and to study the Notch Homolog 1, Translocation-Associated (Notch1), sonic hedgehog (SHH), mammalian target of rapamycin (mTOR), and Wnt/β-catenin signaling pathways in the PAX6 mutant keratocytes.

**METHODS:** Primary human keratocytes were isolated from healthy corneas. Keratocytes were transduced with Cas9 lentiviral particles in order to create cells stably expressing Cas9 nuclease. Lentiviral particles carrying PAX6 sgRNA were transduced into the Cas9 keratocytes creating mutants. Analysis of signaling pathways was assessed by RT-qPCR for gene expression and western blot for protein expression.

**RESULTS:** Human keratocytes stably expressing Cas9 nuclease were created. Keratocytes carrying PAX6 gene mutation were successfully generated. PAX6 mutant keratocytes showed modified expression patterns of extracellular matrix components such as collagens and fibrotic markers. Analysis of the Notch1, SHH, mTOR, and Wnt/β-catenin signaling pathways in the PAX6 mutant keratocytes revealed altered gene and protein expression of the key players involved in these pathways.

**CONCLUSIONS:** A properly functioning PAX6 gene in keratocytes is crucial for the regulation of signaling pathways important for cell fate determination, proliferation, and inflammation. Pax6 mutation in the *in vitro* settings leads to changes in these pathways which resemble those found in corneas of patients with ARK.

## INTRODUCTION

Aniridia is a rare panocular disorder caused by mutations in the PAX6 gene on band p13 of chromosome 11.^1^ It affects the cornea, anterior chamber, iris, lens, retina, macula, and optic nerve resulting in impaired vision from multiple causes including cataract, glaucoma, retinal detachment, and aniridia-related keratopathy (ARK).^2^ The prevalence of the disease in northern American and European populations is estimated to be from 1:64,000 to 1:96,000, with 1:70,000 estimated for Sweden.^3, 4^ It is a complex disorder that is often difficult to manage both medically and surgically.

ARK typically appears after childhood, and both its prevalence and severity of ARK increase with age which lead to recurrent corneal erosions, ulcerations, fibrosis, and opacity, resulting in significant or total loss of vision and decreased quality of life.^1^ ARK is characterized by abnormally differentiated epithelium, impaired epithelial cell adhesion, abnormal healing response, limbal stem cell deficiency and corneal vascular pannus ^2^. The management of ARK depends on its severity and includes preservative-free lubricants in mild ARK, serum drops and amniotic membrane transplants in moderate ARK, and, in severe cases, limbal cell transplant.^5^ Cornea surgery in aniridia patients has a limited prognosis and is therefore avoided as long as possible.^6^

PAX6 is a DNA-binding transcription facto able to directly regulate upstream and downstream transduction pathways.^7, 8^ It has been shown that it maintains human corneal epithelium identity and controls cell differentiation.^9^ PAX6 regulates expression of various cytokeratins,^10^ which form the intermediate filament proteins of the corneal epithelial cells responsible for anchoring the cells to the basal lamina. In aniridia these cytokeratins are downregulated leading to a fragile corneal epithelium.^10, 11^ Moreover, PAX6 regulates expression of adhesion molecules such as β-catenin,^12^ and plays a role in migration of corneal cells and therefore in corneal wound healing.^13, 14^ One of our previous studies ^15^ revealed that the highly conserved Notch1, SHH, mTOR, and Wnt/β-catenin signaling pathways, which are important for cell proliferation, cell renewal and differentiation, and eye development, are involved in aniridia. Using immunohistochemistry we showed augmented detection of the SHH, mTOR, and Wnt/β-catenin signaling pathways, and decreased detection of the Notch1 pathway in corneal tissues of patients with ARK. Notch1 signaling pathway is involved in the regulation of cell-fate both during development and in adult tissue homeostasis and is critical for corneal epithelial repair.^16^ The SHH pathway is associated with regulation of cell differentiation, proliferation and maintenance of tissue polarity. It has been shown to have relevant roles in embryonic development and it is tightly regulated in adult tissues. Aberrant SHH signaling in this pathway has been associated with human cancers.^17^ The serine/threonine kinase mammalian target of rapamycin (mTOR) signaling pathway is considered to be a master growth regulator that integrates various environmental signals^18^ and is crucial in the regulation of several cellular processes including proliferation, growth, transcription, protein synthesis, ribosomal biogenesis and cytoskeletal organization. Altered mTOR signaling has been associated with different eye diseases and embryonic development phenotypes.^19^ Lastly, the Wnt/β-catenin signaling pathway includes several stimulating ligands, including Wnt-5a and Wnt-7a. Wnt-5a is known to be a pro-inflammatory agent,^20^ whereas Wnt-7a is associated with corneal epithelial differentiation control through PAX6. Inhibition of Wnt-7a or PAX6 transforms limbal stem cells into skin-like epithelium.^21^ Target genes of this pathway include fibronectin and vimentin, which are modulators of fibrosis,^22^ and altered Wnt/β-catenin signaling leads frequently to deregulated growth.^23^ These findings are significant in the context of ARK, and are further explored in this paper.

Data collected in human ARK cornea samples ^15^ indicate that these signaling pathways may play key roles in ARK and therefore we sought to develop an in vitro model based on human keratocytes that would allow us to study the relevant molecular changes induced by aniridia/PAX6 gene mutation, how they are regulated, and, in a longer perspective, allow the development of new therapeutic approaches for ARK. In this study we present a new *in vitro* model of ARK in human keratocytes, and study the aforementioned key signaling pathways. Understanding and knowing the key players involved in ARK will possibly allow us to interfere or modulate the signaling pathways, and potentially improve vision of the aniridia patients taking into consideration that there is a window of opportunity between the diagnosis of aniridia in childhood and the clinical onset of ARK during adolescence.

## MATERIALS AND METHODS

### Collection of human corneas

Healthy human corneas were obtained post-mortem from individuals who had chosen, when alive, to donate their corneas for transplantation and research, according to Swedish law, and were kept and processed in the corneal biobank at the University Hospital of Umeå, Sweden. Healthy anterior central lamella from donor’s cornea obtained during corneal endothelial keratoplasties were collected and used for this study which was vetted by the Regional Ethical Review Board in Umeå (2010-373-31M) and performed according to the principles of the Declaration of Helsinki.

### Isolation of primary keratocytes

Isolation and culture of primary keratocytes were performed as previously described.^24^ Briefly, the cornea was cut into small pieces with a scalpel and then digested with 2mg/ml collagenase (Sigma-Aldrich, St. Louis, MO, USA, # C0130) diluted in DMEM/F-12 + GlutaMAX™ medium (Thermo Fisher Scientific, Rockford, IL, USA, # 31330-095) containing 2% fetal bovine serum (FBS; Thermo Fisher Scientific, # 10082-147) and 1% penicillin-streptomycin (Thermo Fisher Scientific, # 15140-122) (DMEM/F-12 2% FBS), overnight at 37°C. Afterwards, samples were centrifuged at 1500 rpm for 5 minutes at room temperature. The pellet was transferred into DMEM/F-12 2% FBS in order to maintain the proper phenotype of quiescent keratocytes. Keratocytes were cultured at 37°C with 5% CO_2_ until they reached confluency. Fresh medium was supplied every second day. 0.05% Trypsin-EDTA (Thermo Fisher Scientific, # 15400-054) was used to detach the cells, which were split in a 1:2 ratio. Cells in passage 3 were used in this study to create keratocytes that stably expressed Cas9 nuclease.

### Generation of keratocytes stably expressing Cas9 nuclease

Invitrogen LentiArray Cas9 Lentivirus (Thermo Fisher Scientific, # A32064) was used to create keratocytes stably expressing Cas9 nuclease. 0.2 × 10^6^ keratocytes were seeded into a 6 well cell culture plates in DMEM/F-12 2% FBS and incubated overnight at 37°C with 5% CO_2_. Next day, medium was replaced with fresh DMEM/F-12 2% FBS containing 8µg/ml of polybrene (hexadimethrine bromide; Sigma-Aldrich, # H9268). Control wells were left without treatment. Invitrogen LentiArray Cas9 Lentivirus (stock virus titer 1 × 10^7^ TU/ml) of MOI (multiplicity of infection) 5 was added dropwise to the cells. The plates were gently swirled and centrifuged at 800 x g for 30 minutes at room temperature. Afterwards, cells were incubated for 2 days at 37°C with 5% CO_2_. Medium was removed and replaced with fresh DMEM/F-12 2% FBS. Cells were incubated for further 2 days at 37°C with 5% CO_2_. In order to select cells positively transduced with the virus, blasticidin (Thermo Fisher Scientific, # A1113903) at a concentration of 17.5 μg/ml was used. Control wells were also treated with blasticidin. Medium containing blasticidin was replaced every 3 days until all cells in the control wells were dead. Blasticidin-resistant cells (the positively transduced) were harvested and serially diluted to a final concentration of 5 cells/ml in DMEM/F-12 2% FBS with blasticidin. 100μl of diluted cells was dispensed into each well of a 96-well plate. Cells were incubated at 37°C with 5% CO_2_ with fresh medium containing blasticidin supplied every 3 days. After 10 days of incubation, colonies that were derived from a single cell were transferred into separate wells of a 24-well plate in medium with blasticidin. Every 2-5 days each clonal population was transferred to a larger plate and later on T-25 and T-75 flasks. After reaching confluency, cells (Cas9 keratocytes) were cryopreserved and characterized for expression of keratocyte markers, and Cas9 protein expression.

### Generation of PAX6 mutants

Invitrogen LentiArray CRISPR PAX6 gRNA Lentivirus (Thermo Fisher Scientific; CRISPR620221_LV) which is a pre-designed, and pre-packaged gRNA lentivirus, was used to create PAX6 mutants in keratocytes stably expressing Cas9 nuclease. This virus targets the CCGAGACAGATTACTGTCCG DNA sequence of the PAX6 gene which is located on Chr.11: 31801590 – 31801612 on GRCh38. 0.2 × 10^6^ keratocytes were seeded into 6-well plates in DMEM/F-12 2% FBS and incubated overnight at 37°C with 5% CO_2_. Next day, medium was replaced with fresh DMEM/F-12 2% FBS containing 8μg/ml of polybrene. Control wells were left without treatment. Invitrogen LentiArray CRISPR PAX6 gRNA Lentivirus (stock virus titer 1 × 10^6^ TU/ml) of MOI 5 was added dropwise to the cells. The plates were gently swirled and centrifuged at 800 × g for 30 minutes at room temperature. Afterwards, cells were incubated for 2 days at 37°C with 5% CO_2_. Medium was removed and replaced with fresh DMEM/F-12 2% FBS. Cells were incubated for further 2 days at 37°C with 5% CO_2_. In order to select cells positively transduced with the virus, puromycin (Thermo Fisher Scientific, # A1113803) at a concentration of 5μg/ml was used. Control wells were also treated with puromycin. Medium containing puromycin was replaced every 3 days until all cells in the control wells were dead. After positive selection, cells were collected and analyzed.

### RT-qPCR

After positive selection of the PAX6 mutants, cells were lysed, and mRNA was extracted using RNA extraction kit (Qiagen, Venlo, Netherlands, # 74106) according to manufacturer’s instructions. Shortly, a high-capacity cDNA reverse transcription kit (Thermo Fisher Scientific, # 4368813) was used to reverse transcribe 1000ng of RNA into cDNA. Collagen I (COL1A1), collagen III (COL3A1), collagen IV (COL4A1), collagen V (COL5A1), lumican (LUM), fibronectin (FN), α-smooth muscle actin (ACTA2), keratocan (KER), CD34, aldehyde dehydrogenase 1 family, member A1 (ALDH1A1), aldehyde dehydrogenase 3 family, member A1 (ALDH3A1), NOTCH1, hairy and enhancer of split-1 (HES1), SHH, mTOR, ribosomal protein S6 (RPS6), WNT5A, β-catenin (CTNNB1) probes were used in order to determine gene expression (Thermo Fisher Scientific). Samples were run in duplicates in ViiA™ 7 Real-Time PCR System (Thermo Fisher Scientific). 18S and β-actin served as endogenous controls (Thermo Fisher Scientific; # 4333760F and # 4352935E, respectively). Analysis was performed with ViiA™ 7 Software (Thermo Fisher Scientific).

### Western blot

After positive selection of the PAX6 mutants, cells were freeze/thawed 3 times and further lysed with RIPA (radioimmunoprecipitation) lysis buffer (Thermo Fisher Scientific, # 89901) supplemented with protease and phosphatase inhibitor cocktail (Thermo Fisher Scientific, # 78446). Total protein concentration was assessed with Bradford assay (Bio-Rad, Hercules, CA, USA, # 5000006). SDS-polyacrylamide gels (Bio-Rad) were used to separate samples. Next, samples were transferred to PVDF membranes (GE Healthcare, Little Chalfont, UK, # GEHERPN303F) and blocked with 5% (w/v) bovine serum albumin (BSA; Sigma-Aldrich, # A9647) or 5% (w/v) non-fat dry milk in TRIS-buffered saline (TBS) containing 0.1% Tween-20 (TBS-T; VWR, Radnor, PA, USA, # 28829.183) for one hour at room temperature and incubated with primary antibodies overnight at 4°C. The following primary antibodies were used: anti-CRISPR-Cas9 (Abcam, Cambridge, UK, # ab191468), anti-NOTCH1 (Abcam, # ab52627), anti-Wnt5 (Abcam, # ab72583), anti-NUMB (Cell Signaling, Danvers, MA, USA, # 2756), anti-p-NUMB (Cell Signaling, # 9878), anti-β-catenin (Cell Signaling, # 8480), anti-Hes1 (Genetex, Irvine, CA, USA, # GTX108356), RPS6 (Genetex, # GTX60800), and anti-β-actin antibody (Cell Signaling, # 4967). Afterwards, membranes were washed in TBS-T and incubated with anti-rabbit IgG HRP-linked secondary antibody (Cell Signaling, # 7074), or anti-mouse IgG HRP-linked secondary antibody (Cell Signaling, # 7076) for one hour at room temperature. Images were taken by Odyssey^®^ Fc imaging system (LI-COR, Lincoln, NE, USA).

### Statistical analysis

All experiments were performed in triplicates. Data are presented as mean ± SD. Statistical analysis was performed with one-way ANOVA with Tukey post hoc test or unpaired t-test. A p-value of < 0.05 was considered statistically significant.

## RESULTS

### Cas9 transduced keratocytes stably express Cas9 nuclease and continue to express keratocytes markers

The first step for establishing the ARK *in vitro* model was to create keratocytes that would stably express the Cas9 nuclease. To achieve that, keratocytes were transduced with lentiviral particles carrying the Cas9 gene. Later on, the positive cells were selected with blasticidin and clonally expanded. We picked 5 clones that were expanded and analyzed for Cas9 expression. Western blot analysis showed that all 5 clones expressed the Cas9 nuclease (Fig. 1A). Moreover, we analyzed gene expression of typical keratocyte markers in the Cas9 keratocytes (Fig. 1B) and expression of keratocan, lumican, and ALDH3 was downregulated in the Cas9 keratocytes. Expression of CD34, and ALDH1 was unaffected by the Cas9 lentiviral transduction. Additionally, we also investigated gene expression of α-SMA, a marker for contractile myofibroblasts, and we found that the transduction of the Cas9 lentiviral particles upregulated its gene expression (Fig. 1B).

**Figure 1.**
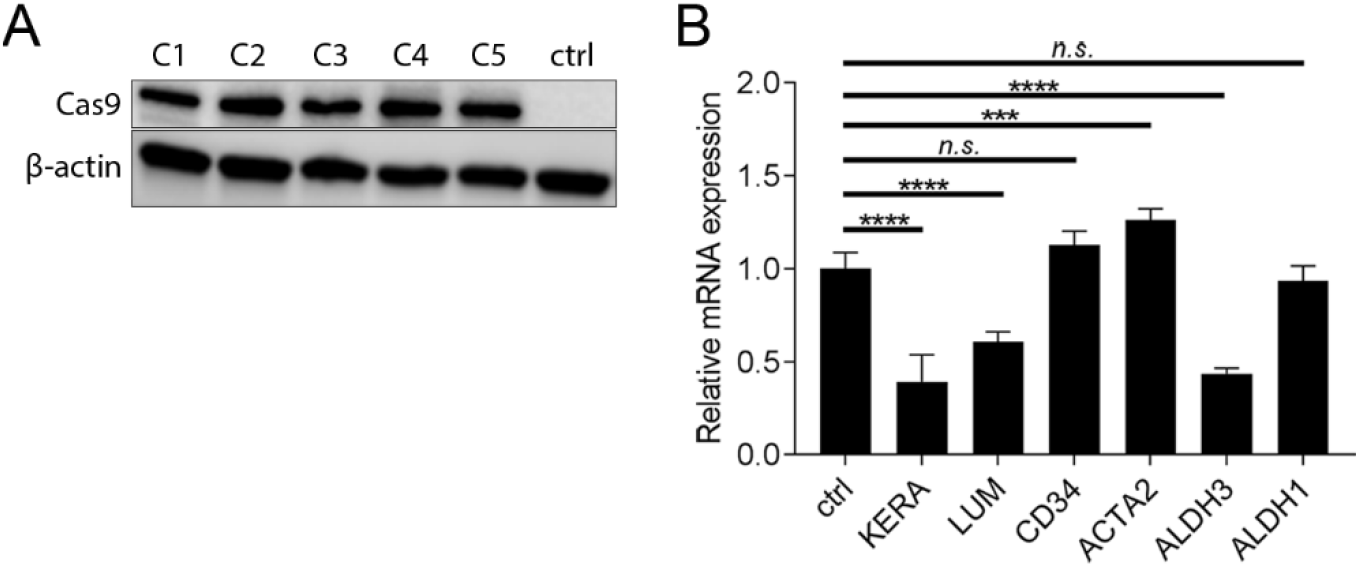
Cas9 transduced keratocytes stably express Cas9 nuclease and continue to express keratocytes markers. (**A**) 0.2 × 10^6^ keratocytes were seeded into 6 well cell culture plates in DMEM/F-12 2% FBS. LentiArray Cas9 Lentivirus of MOI 5 was added dropwise to the cells. Control wells were left without treatment. Cells were incubated for 2 days at 37°C with 5% CO_2_. Positively transduced cells were selected by blasticidin (17.5 μg/ml). Medium containing blasticidin was replaced every 3 days until cells in the control wells were all dead. Blasticidin-resistant cells were harvested and serially diluted to a final concentration of 5 cells/ml in DMEM/F-12 2% FBS with blasticidin and clonally expanded. Cells originating from 5 different clones showed stable expression of Cas9 nuclease (160 kDa) as assessed by western blot. β actin (45kDa) served as loading control. (**B**) Keratocytes stably expressing Cas9 nuclease were lysed and total RNA was extracted and reversed transcribed into cDNA. Gene expression of keratocan (KERA), lumican (LUM), CD34, α-SMA (ACTA2), ALDH3, and ALDH1 was determined, and compared to untransduced keratocytes. No significant change in CD34 and ALDH1 expression was observed. Gene expression of ACTA2 increased, with expression of KERA, LUM, and ALDH3 significantly decreased. Values are means ± SD. n.s. (not significant); ***p<0.001, ****p<0.0001.

### PAX6 mutant keratocytes show altered gene expression of collagens and fibrotic markers

The main role of keratocytes is to produce components of the extracellular matrix (ECM) such as collagens. Therefore, we were interested whether mutation of the PAX6 gene in keratocytes would affect the ECM production. We found that gene expression of the two main types of collagens found in the cornea, collagen I and collagen V, was downregulated in PAX6 mutant keratocytes (Fig. 2A). Expression of collagen III, which is expressed in the cornea during wound healing, was downregulated in the PAX6 mutants (Fig.2B). Collagen IV, usually present in the epithelial basement layer, was overexpressed in the PAX6 mutant keratocytes (Fig. 2B). Additionally, the PAX6 mutant keratocytes showed downregulation of two fibrotic markers genes, α-SMA and fibronectin (Fig. 2C).

**Figure 2.**
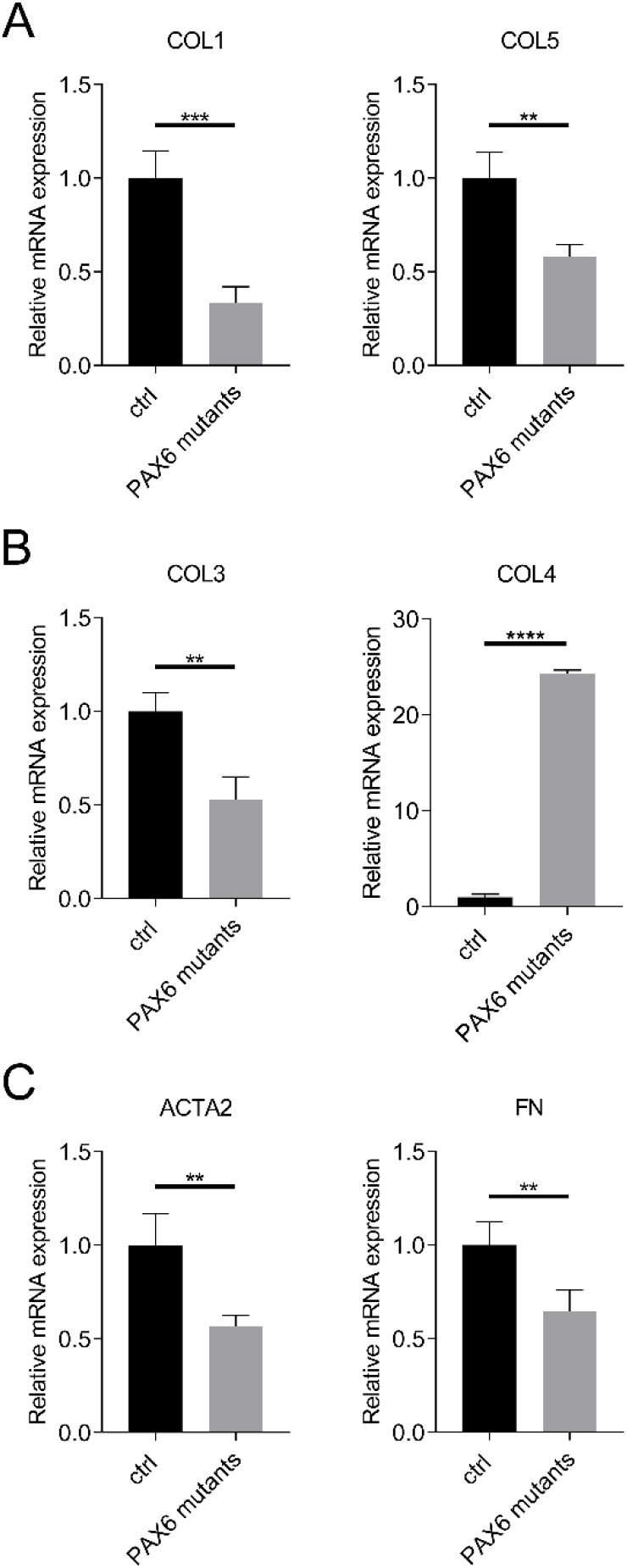
PAX6 mutant keratocytes show altered gene expression of collagens and fibrotic markers. PAX6 mutant keratocytes were lysed, total RNA was extracted and reversed transcribed into cDNA. (**A**) Gene expression of collagen I (COL1) and collagen V (COL5) was decreased in PAX6 mutant keratocytes. (**B**) Gene expression of collagen III (COL3) was decreased, and gene expression of collagen IV (COL4) was increased in PAX6 mutant keratocytes. (**C**) Gene expression of ACTA2 and fibronectin (FN) was decreased in PAX6 mutant keratocytes. Gene expression in PAX6 mutant keratocytes was compared to untransduced Cas9 keratocytes. Values are means ± SD. **p<0.01, ***p<0.001, ***p<0.0001.

### Notch1 and SHH pathways are altered in PAX6 mutant keratocytes

PAX6 mutation in keratocytes had no effect on the expression of the NOTCH1 gene. However, as analyzed by western blot, the PAX6 mutation resulted in decreased production of the Notch1 protein (Fig. 3A). Expression of NUMB, the Notch1 inhibitor, was decreased in PAX6 mutants. Both the unphosphorylated and phosphorylated forms of Numb were downregulated in PAX6 mutant keratocytes (Fig. 3B). Gene expression of HES1, the downstream target of NOTCH1, was downregulated in PAX6 mutants, however the production of Hes1 protein was upregulated in the mutants (Fig. 3C). Moreover, gene expression of SHH was increased in the PAX6 mutants (Fig. 3D).

**Figure 3.**
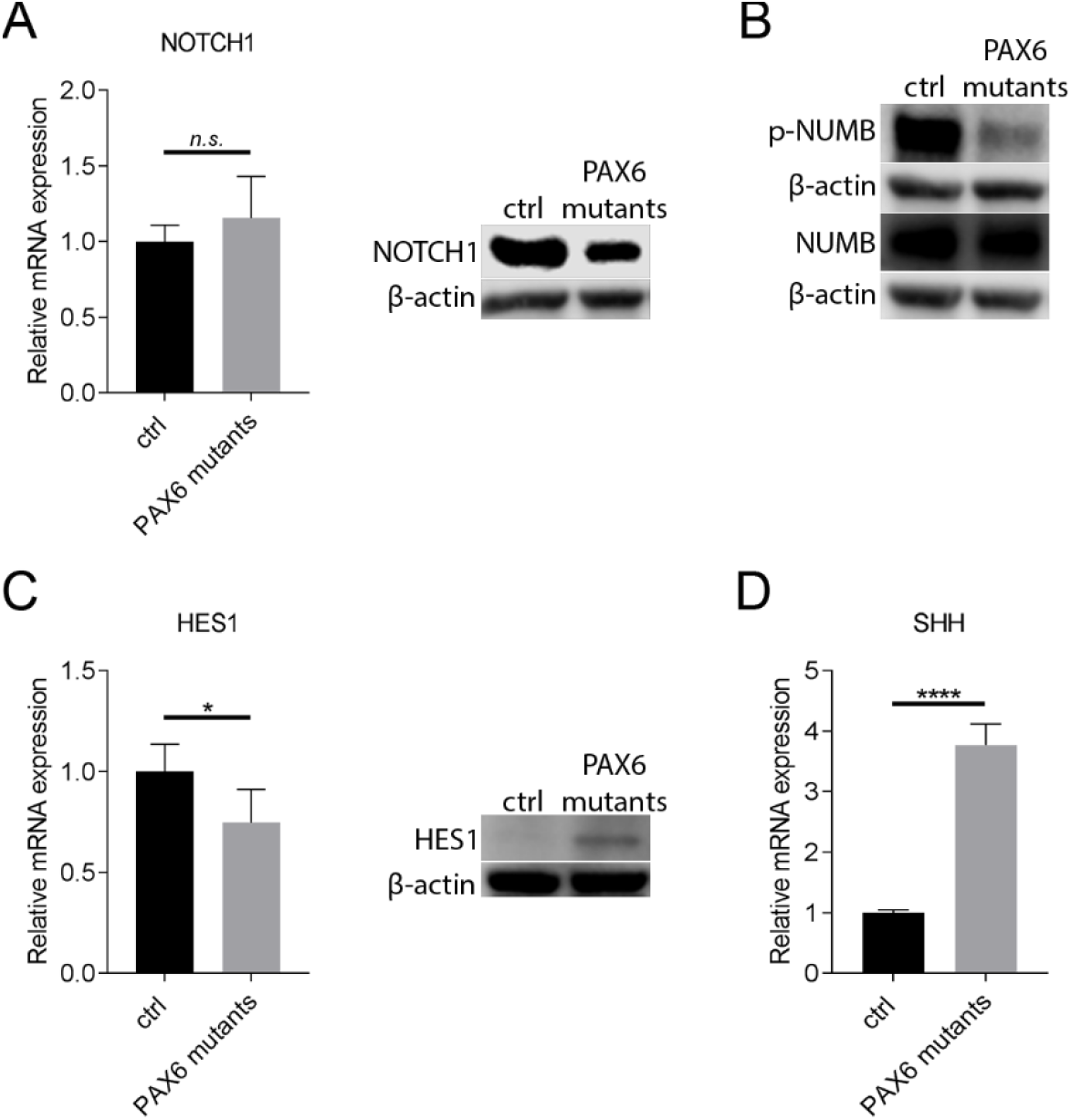
NOTCH1 and SHH pathways are altered in PAX6 mutant keratocytes. (**A**) PAX6 mutant keratocytes were lysed, total protein and total RNA was extracted. RNA was further reversed transcribed into cDNA. Gene expression of NOTCH1 was not affected in PAX6 mutant keratocytes. Western blot analysis showed that NOTCH1 (125 kDa) expression is downregulated in PAX6 mutant keratocytes. β-actin (45kDa) served as loading control. (**B**) PAX6 mutant keratocytes were lysed and total protein was extracted. Expression of both p-NUMB (Ser276) (78 kDa), and NUMB (78 kDa) were decreased in PAX6 mutant keratocytes. β-actin (45kDa) served as loading control. (**C**) PAX6 mutant keratocytes were lysed, total protein and total RNA was extracted. RNA was further reversed transcribed into cDNA. Gene expression of HES1 was downregulated in PAX6 mutant keratocytes, however expression of HES1 protein (30 kDa) was increased. β-actin (45kDa) served as loading control. (**D**) PAX6 mutant keratocytes were lysed, total RNA was extracted and reversed transcribed into cDNA. Gene expression of Sonic Hedgehog (SHH) was upregulated in PAX6 mutant keratocytes. Gene expression in PAX6 mutant keratocytes was compared to untransduced Cas9 keratocytes. Values are means ± SD. n.s. (not significant); *p<0.05, ****p<0.0001.

### mTOR signaling is altered in PAX6 mutant keratocytes

The mutation of PAX6 resulted in increased expression of the mTOR gene (Fig. 4A). PAX6 mutant keratocytes had downregulated gene expression of RPS6, a downstream target of mTOR. However, rpS6 protein levels were increased in the PAX6 mutant keratocytes (Fig. 4B).

**Figure 4.**
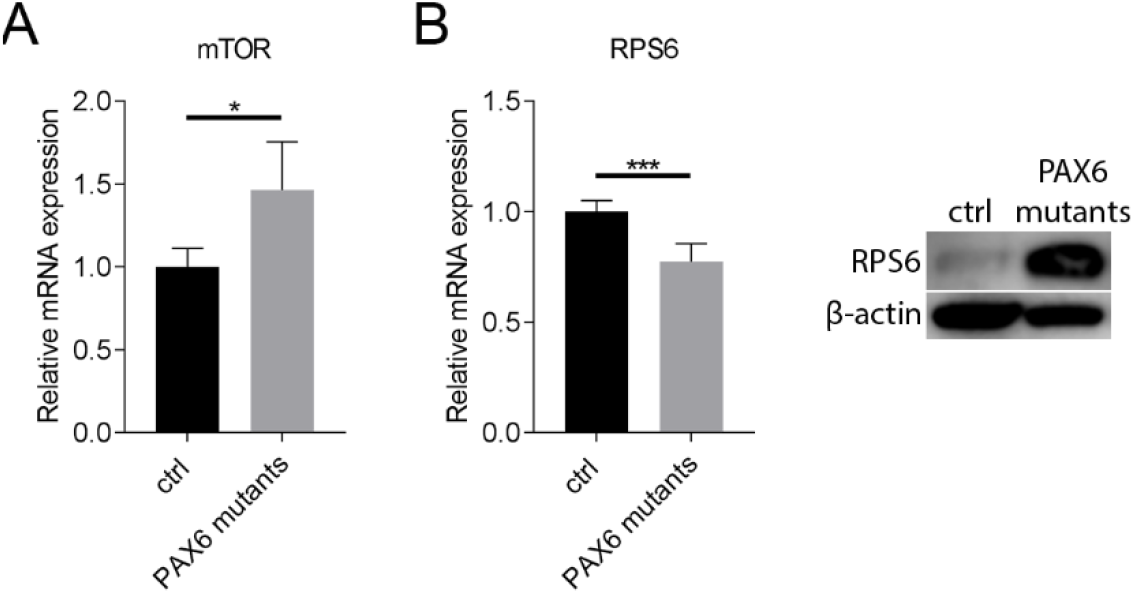
mTOR signaling is altered in PAX6 mutant keratocytes. (**A**) PAX6 mutant keratocytes were lysed, total RNA was extracted and reversed transcribed into cDNA. Gene expression of mTOR was upregulated in PAX6 mutant keratocytes. (**B**) PAX6 mutant keratocytes were lysed, total protein and total RNA was extracted. RNA was further reversed transcribed into cDNA. Gene expression of RPS6 was downregulated in PAX6 mutant keratocytes, however expression of RPS6 protein was increased. β-actin (45kDa) served as loading control. Gene expression in PAX6 mutant keratocytes was compared to untransduced Cas9 keratocytes. Values are means ± SD. *p<0.05, ***p<0.001.

**Figure 5.**
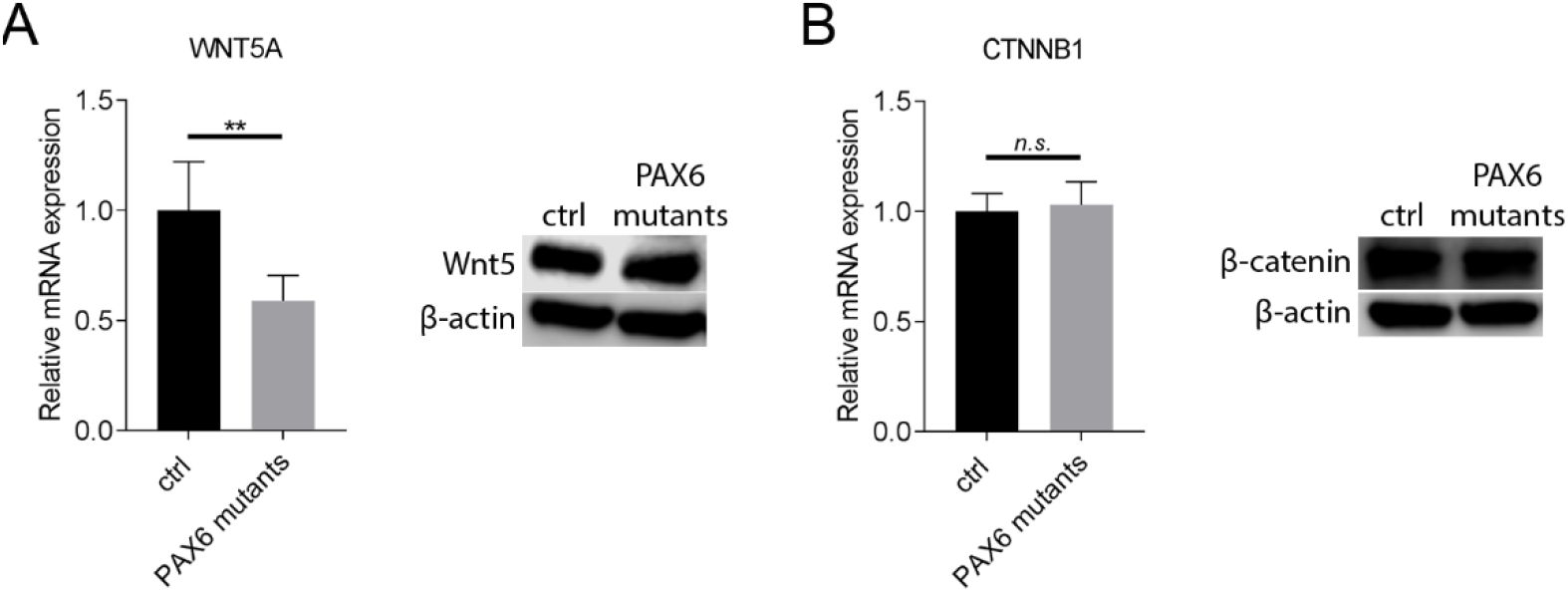
Wnt signaling is altered in PAX6 mutant keratocytes. (**A**) PAX6 mutant keratocytes were lysed, total protein and total RNA was extracted. RNA was further reversed transcribed into cDNA. Gene expression of WNT5A was downregulated in PAX6 mutant keratocytes. Expression of WNT5 protein was unaltered in PAX6 mutants. β-actin (45kDa) served as loading control. (**B**) PAX6 mutant keratocytes were lysed, total protein and total RNA was extracted. RNA was further reversed transcribed into cDNA. Neither gene expression nor protein expression of β-catenin (CTNNB1) was affected in PAX6 mutant keratocytes. β-actin (45kDa) served as loading control. Gene expression in PAX6 mutant keratocytes was compared to untransduced Cas9 keratocytes. Values are means ± SD. n.s. (not significant); **p<0.01.

### Wnt/β-catenin signaling is altered in PAX6 mutant keratocytes

PAX6 mutant keratocytes showed decreased gene expression of the WNT5A gene, however, the levels of Wnt5A protein remained unchanged. The gene expression of β-catenin, the downstream target in Wnt/β-catenin, signaling pathway was unaltered in PAX6 mutant keratocytes but its protein levels were decreased.

## DISCUSSION

In this study, we introduce for the first time an *in vitro* ARK model of CRISPR/Cas9 engineered human keratocytes, and we are able to show that crucial signaling pathways, such as Notch1, SHH, mTOR, and Wnt/β-catenin pathways, responsible for cell proliferation, cell renewal and differentiation and eye development, resemble the altered signaling pattern found in human corneal samples of patients with ARK.^15, 25^ Aniridia and the resulting ARK are difficult to manage both medically and surgically.^5^ A number of surgical procedures is available but, unfortunately, their outcome is poor.^6^ Because of poor healing response and limbal stem cell deficiency, penetrating keratoplasty has a poor prognosis in aniridia corneas.^26^ Another treatment option available for ARK patients is keratolimbal allograft transplantation (KLAL) but the graft survival rate decreases dramatically over a 2-year period and patients need to take aggressive systemic immunosuppressive medications.^27^ Additionally, there is a recurrence of ARK symptoms after surgery, due to the impaired corneal healing and fibrosis.^1^ Therefore, despite the increasing knowledge about the genetics and pathology of ARK, there is still lack of effective treatment approaches. Moreover, due to the limited prognosis of surgery, it is very difficult to acquire corneal samples form ARK patients. Consequently, it is difficult to study the role of PAX6 in the pathophysiology of ARK, specifically regarding cell signaling pathways, which could possibly lead to the development of new therapeutic approaches. Thus, an *in vitro* ARK model was needed and we established an *in vitro* model of aniridia/ARK using CRISPR/Cas9 engineered human keratocytes in which the PAX6 gene is mutated to mimic the disease, in order to identify key molecules and molecular pathways responsible for the ARK, and, as an end point in future studies, to facilitate screening for new potential therapeutic approaches. In order to establish the model we used primary keratocytes isolated from healthy anterior central lamella of the cornea, as we have reported earlier ^25^ that the structural changes in ARK patients are mostly present in the anterior part of the cornea. First, through transduction of lentiviral particles, and selection with blasticidin, we established keratocytes stably expressing Cas9 polymerase. These cells expressed common keratocyte markers such as keratocan, lumican, CD34, ALDH3, and ALDH1. Slightly increased expression of α-SMA was observed, however, taken into consideration that the cells expressed keratocyte markers, we concluded that they were not in an activated state. Lentiviral particles containing PAX6 gRNA were transduced into the Cas9 keratocytes with an efficiency of ~80% (data not shown). We were able to select the PAX6 mutants with puromycin. Moreover, we confirmed that the PAX6 transduced keratocytes did not express PAX6 gene through RT-PCR (data not shown). However, as PAX6 expression levels in healthy keratocytes are very low, we believe that there is a need for further confirmation of the PAX6 mutation in our model. DNA sequencing is underway in order to confirm and further assess the mutation.

Previous studies in our lab ^25^ showed that naïve ARK corneas (corneal buttons from patients with advanced ARK submitted to penetrating keratoplasty for the first time), and transplanted corneas (from patients who were re-transplanted due to failure of a former centered or decentered transplant with a cornea from a healthy donor) had altered expression of extracellular matrix components such as collagen I and collagen IV, and showed increased expression of fibrotic markers such as α-SMA and fibronectin. The results from our *in vitro* model show similar changes: lowered expression of the two major types of collagens present in the cornea (I and V),^28^ decreased expression of collagen III, which is present in the cornea during pathological and wound healing processes.^29^ A striking increase in the expression of collagen IV, a network-forming collagen in basement membranes ^30^ was also consistent with the results from ARK and transplanted corneas (REF). Interestingly, we found that gene expression α-SMA and fibronectin was decreased in the *in vitro* model, which contradicts the results found in naïve and transplanted ARK patients. However, in order to compare the *in vitro* results with those from the patient samples, we think that further analysis of α-SMA protein expression and secreted/extracellular levels of fibronectin in our *in vitro* model is needed as it will better relate to the *in vivo* findings. The main aim of this study was to investigate whether the PAX6 mutation created *in vitro* will affect the highly conserved signaling pathways: Notch1, SHH, mTOR, and Wnt/β-catenin pathways, shown to be affected in our previous studies based on samples of ARK patients.^15^ In line with the patients’ findings, the Notch1 protein expression was decreased in the ARK *in vitro* model, however, its gene expression was unaffected by the PAX6 mutation. The *in vitro* model showed that NUMB, a key regulator of cell fate, and Notch1 inhibitor,^31^ was decreased in both phosphorylated and unphosphorylated state, which is the opposite of what was found in ARK patients’ samples. Perhaps, in the *in vitro* system, the control of Notch1 signaling by Numb is abrogated by ubiquitination and proteasomal degradation of Numb.^32^ Hes1 is a classical downstream target of Notch1 signaling ^33^, and we have previously shown that its expression is increased in naïve and transplanted ARK patients. In the *in vitro* model, Hes1 protein was increased, however the gene expression was lower than in healthy keratocytes, suggesting posttranscriptional, posttranslational modifications, or perhaps an accumulation of Hes1 protein in the cells. Additionally, Hes1 is induced by SHH, and this induction is required for SHH mediated cell proliferation.^33^ We were able to show that gene expression of SHH in the PAX6 mutants was increased, suggesting its role in the induction of Hes1 in ARK. The results from patients’ corneas also showed that mTOR pathway is altered in ARK. Interaction between mTOR and its downstream effector rpS6 controls fundamental cellular processes such as transcription, translation, and cell proliferation.^34^ The PAX6 mutant keratocytes presented increased expression of mTOR but lower expression of RPS6 genes. However, the expression of rpS6 protein was greatly increased in the PAX6 mutant keratocytes, again, suggesting posttranscriptional, posttranslational modifications, or accumulation of the rpS6 protein in the cells. The Wnt/β-catenin pathway is an evolutionarily conserved pathway that regulates cell fate determination, cell migration and organogenesis during embryonic development, among many other functions.^35^ The naïve and transplant ARK patients had increased amounts of Wnt5, a noncanonical Wnt family member. However, expression of the WNT5 gene in the ARK *in vitro* model was decreased with no difference in protein expression between healthy and PAX6 mutant keratocytes. Moreover, in the ARK *in vitro* model, we found no differences in expression of β-catenin, a key nuclear effector of canonical Wnt signaling in the nucleus,^36^ between healthy keratocytes and PAX6 mutant keratocytes on both gene and protein levels. In contrast, in the naïve and transplanted ARK patients, β-catenin was observed in epithelial cells, blood vessels, and in the subepithelial pannus. This differences might be a consequence of the existence of different stimuli in the setting of chronic inflammation in ARK corneas, that are absent in the *in vitro* model, or arise from the influence of the corneal epithelium in the stroma in the absence of a proper basement membrane between the two layers in the ARK corneas.

In summary, the altered signaling pathways previously found in ARK patients are present in the new ARK *in vitro* model of PAX6 mutant keratocytes. As aniridia patients are usually diagnosed several years prior to the development of ARK, there is a window of opportunity for novel therapeutic approaches aimed at maintaining epithelial integrity and corneal transparency. We think that our ARK *in vitro* model has great potential value for studying the molecular mechanisms of ARK pathophysiology and for screening of possible future therapeutic approaches.

## ACKNOWLEDGMENTS

The authors thank Dr. Maria Brohlin, and Ms. Randi Elstad for help in providing the donated corneas from the biobank.

## Notes

### Competing Interest Statement

The authors have declared no competing interest.

